# Leveraging allele-specific expression to refine fine-mapping for eQTL studies

**DOI:** 10.1101/257279

**Authors:** Jennifer Zou, Farhad Hormozdiari, Brandon Jew, Jason Ernst, Jae Hoon Sul, Eleazar Eskin

## Abstract

Many disease risk loci identified in genome-wide association studies are present in non-coding regions of the genome. It is hypothesized that these variants affect complex traits by acting as expression quantitative trait loci (eQTLs) that influence expression of nearby genes. This indicates that many causal variants for complex traits are likely to be causal variants for gene expression. Hence, identifying causal variants for gene expression is important for elucidating the genetic basis of not only gene expression but also complex traits. However, detecting causal variants is challenging due to complex genetic correlation among variants known as linkage disequilibrium (LD) and the presence of multiple causal variants within a locus. Although several fine-mapping approaches have been developed to overcome these challenges, they may produce large sets of putative causal variants when true causal variants are in high LD with many non-causal variants. In eQTL studies, there is an additional source of information that can be used to improve fine-mapping called allele-specific expression (ASE) that measures imbalance in gene expression due to different alleles. In this work, we develop a novel statistical method that leverages both ASE and eQTL information to detect causal variants that regulate gene expression. We illustrate through simulations and application to the Genotype-Tissue Expression (GTEx) dataset that our method identifies the true causal variants with higher specificity than an approach that uses only eQTL information. In the GTEx dataset, our method achieves the median reduction rate of 11% in the number of putative causal variants.

## 1 Introduction

Understanding the regulation of gene expression by genetic variants is essential for identifying the biological mechanisms of gene expression and complex traits. In expression quantitative trait loci (eQTL) studies, a statistical test is performed to find genetic variants called eQTLs that are significantly correlated with gene expression. Identifying eQTLs is critical in genetic studies not only because they influence gene expression but also because they are enriched in disease risk loci [1, 2, 3, 4, 5]. In recent years, many studies have identified eQTLs in different organisms and tissues [6, 7, 8, 9, 10].

Once eQTLs are identified, the next step is to isolate the causal variants that influence gene expression. In genetics, causal variants are variants that are responsible for the observed peak of association. Not all eQTLs are causal, and identifying causal variants from several candidate variants is called fine-mapping. Fine-mapping in eQTL studies has two main challenges. The first challenge is the complex LD structure present in human genome. If a region of the genome contains many genetic variants that are highly correlated (or “in high LD”) with each other, non-causal genetic variants close to a causal variant appear to be correlated with gene expression [11, 12, 13]. The second challenge is that there may be multiple causal variants in a region [14, 15], increasing the complexity of fine-mapping algorithms. A few fine-mapping approaches have been developed to address these challenges [16, 17, 18, 19, 20, 21, 22]. These methods attempt to calculate a posterior probability for each variant and select a set of variants called a “causal set” that contains all causal variants with high probability (e.g., 95%), while minimizing the number of variants in the causal set. Minimizing the causal set size reduces the number of variants that need to be validated using biological assays. Although these previous methods are accurate in including all true causal variants in the causal set, they may also include many non-causal variants in regions with high LD, which increases the cost of biological validation.

In addition to eQTL data, there is another source of information called allele-specific expression (ASE) that we can utilize to identify genetic variants that regulate gene expression [23, 24, 25, 26, 27, 28]. An individual has ASE for a specific gene if the amount of gene expression from one haplotype is greater or smaller than that from the other haplotype (”allelic imbalance”) [29, 30]. Cis-regulatory variants that cause ASE may be identified by finding association between the ASE status and heterozygous status of a variant across individuals [31, 26]. For example, if an individual has a heterozygous genotype at a causal variant, we expect the amount of gene expression from two alleles to be different. On the other hand, if an individual has a homozygous genotype at a causal variant, we expect the amount of gene expression to be balanced among two alleles (Figure 1A). Because causal variants for ASE change expression levels of a gene in an allele-specific manner, they provide insight into the regulation of gene expression. It has also been shown that ASE is enriched in eQTLs [32], indicating that genetic variants causing changes in total gene expression are also likely to cause ASE. Several studies have combined eQTL and ASE data to improve the power of eQTL studies, which leads to identification of additional eQTLs [33, 34, 35]. However, these methods often focus on improving power rather than reducing the number of putative causal variants and fail to take multiple causal variants into account.

**Figure 1:**
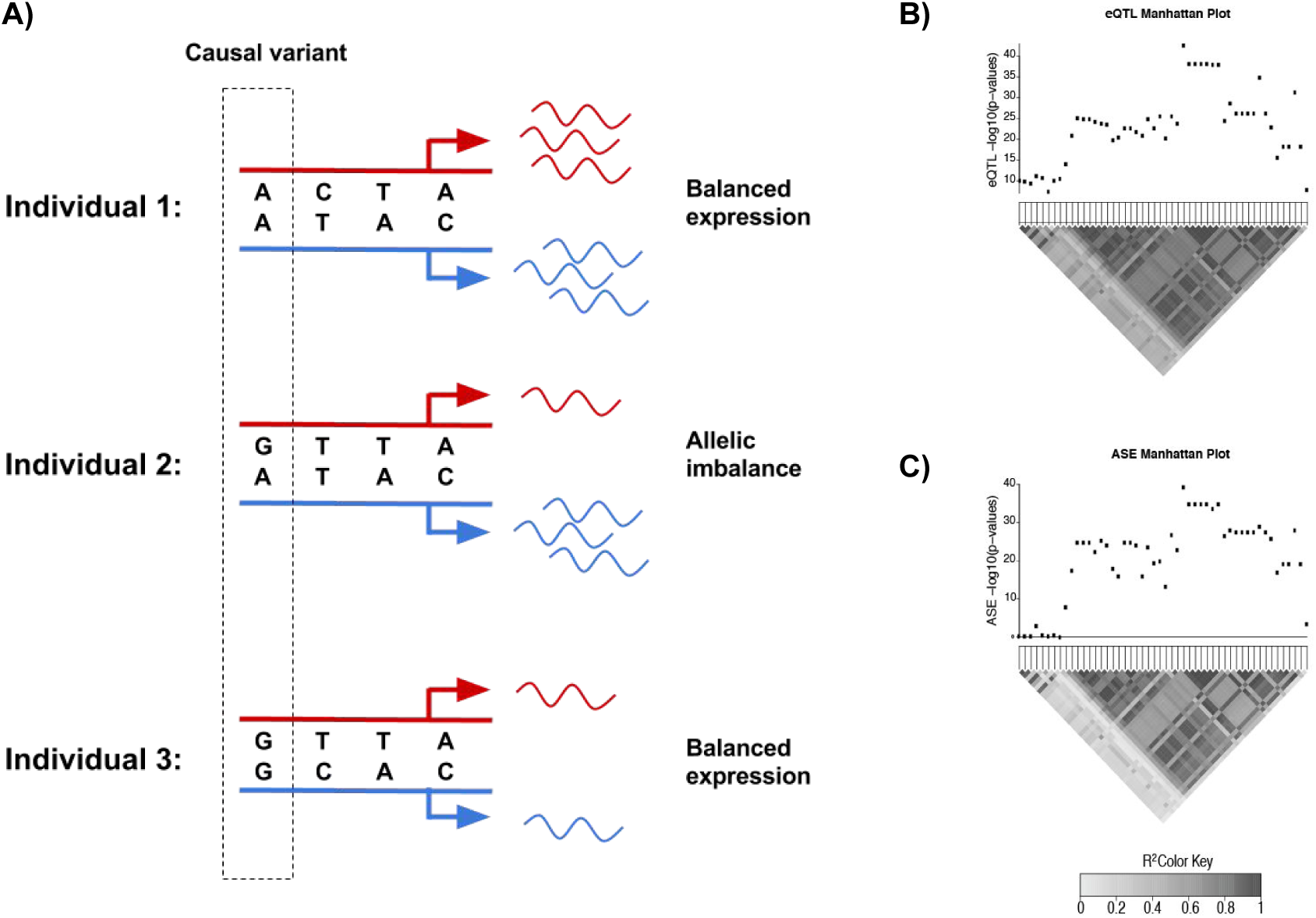
A) Individuals 1 and 3 have balanced expression, and individual 2 has ASE. The first variant fits this pattern of ASE status across individuals the best among the three variants shown. B and C) Manhattan plots for gene ENSG00000115705 showing *-log_10_* p-values for eQTL and ASE statistics respectively. The heatmap below the eQTL plot shows pairwise correlation of genotypes, and the heatmap below the ASE plot shows pairwise correlation of heterozygosity.

In this paper, we propose a method that combines both eQTL and ASE information of genetic variants to improve fine-mapping in eQTL studies. We first compute an eQTL statistic for total expression and an ASE statistic for allele-specific expression, and we then aggregate the two statistics utilizing meta-analysis. The meta-analysis statistic is then incorporated into a fine-mapping approach for GWAS called CAVIAR [18], which takes into account the LD structure among genetic variants and incorporates multiple causal variants. Our simulation results show that our method identifies causal variants with the correct true positive rate or sensitivity, and more importantly, it generates causal sets that are noticeably smaller than sets generated using only the eQTL statistics. This means that our approach includes fewer non-casual variants, yielding higher specificity. We apply our method to the RNA-seq data from ten tissues in the Genotype-Tissue Expression (GTEx) dataset [8, 10] and find that the causal set size is also reduced when compared to an approach that only uses eQTL statistics.

## 2. Results

### 2.1 Method overview

The main goal of fine-mapping methods is to identify a small set of variants that include all true causal variants with high probability. A naive approach would be to conduct an eQTL mapping and to use these statistics as input to fine-mapping algorithms for GWAS such as CAVIAR [18]. Our method improves this fine-mapping algorithm by including ASE signals derived from RNA-seq data. In ASE association mapping, we are interested in finding genetic variants that are associated with a significant difference in expression between two haplotypes. This is different from eQTL mapping, which identifies genetic variants associated with a significant difference in total expression. Thus, the ASE analysis may provide additional information on which variants are causal by using the expression difference between the two haplotypes, which is not captured in the eQTL mapping.

In terms of a statistic for each analysis, an eQTL analysis computes a statistic that measures correlation between total expression levels of a gene and genotypes of a variant. On the other hand, an ASE analysis computes a statistic that measures correlation between ASE status (e.g., 0 for balanced expression and 1 for allelic imbalance) and heterozygous status of a variant (e.g., 0 for homozygous genotype and 1 for heterozygous genotype). In Figure 1A, individual 2 who has a heterozygous genotype at the causal variant has allelic imbalance, while other individuals with homozygous genotypes have balanced expression. The causal variant would have a higher ASE statistic than any of the other non-causal variants if it truly affects the gene expression. We compute the ASE statistic as discussed in the Methods section.

Because ASE statistics measure cis-regulatory strength of variants, they can be used to improve fine-mapping. For example, a certain gene may contain many highly correlated genetic variants with low p-values using eQTL mapping (Figure 1B), which makes it difficult to pinpoint causal variants. In this case, ASE statistics may provide more information on which of those variants are more likely to be causal (Figure 1C). Hence, by combining eQTL and ASE statistics, we can detect the true causal variants more accurately.

Given these two different statistics, we assume that causal variants in the eQTL analysis also cause changes in ASE status. This assumption is likely to be valid because an allele that increases the total expression level of a gene causes an individual with a heterozygous genotype to have a different relative contribution from two alleles to the total expression level (or ASE). With this assumption, we combine eQTL and ASE summary statistics of variants in close proximity to a gene by performing meta-analysis on the two statistics. Meta-analysis is a popular statistical approach that combines results of multiple studies to increase statistical power. In GWAS, meta-analysis has discovered many associations that were not identified by each individual study. We perform meta-analysis between the ASE and eQTL statistics to better identify causal variants that influence gene expression.

To improve identification of causal variants from these statistics, we extend the previous fine-mapping framework for GWAS called CAVIAR[18]. CAVIAR takes as input a set of summary statistics for variants in a locus and a pairwise correlation matrix between these variants calculated from genotype data. Using this data, CAVIAR identifies a subset of variants that contains all causal variants with probability *ρ*. We refer to this set as the *ρ* causal set, and this can be interpreted as a confidence interval on the set. We extend the CAVIAR framework to incorporate both eQTL and ASE data as discussed in the Methods section. By integrating these two types of data, our method attempts to reduce the size of the causal set, increasing specificity compared to the traditional CAVIAR approach that uses only eQTL data.

### 2.2 Meta-analysis achieves correct recall rate

We generate simulated data to assess the performance of our approach. In this simulation, we use real genotypes from the whole blood tissue in the GTEx dataset that contains 325 samples. For each gene, we identify the top 50 genetic variants with the highest eQTL statistics. We use these top 50 genetic variants to calculate pairwise correlation matrices as described in the Methods section. We then choose one causal variant randomly from the 50 variants and assign an effect size to this variant such that we have 50% power to detect its association. Given this effect size for the causal variant, we generate an eQTL statistic for the causal variant and eQTL statistics for the remaining variants by using their genotype correlation (LD) to the causal variant and by assuming that the statistics follow the multivariate normal distribution (MVN). Using the same effect size and implanted causal variant, we generate an ASE statistic for the causal variant and ASE statistics for the rest of the variants by using their heterozygosity correlation to the causal variant and the same MVN assumption. We combine these simulated statistics into a meta-analyzed statistic (”meta-statistic”), which we use as input to our fine-mapping framework. For comparison, we apply our approach using only the simulated eQTL statistics. We also generate simulations where the number of causal variants is 2 and 3 and the power to detect a causal variant is 80%. We use 1,000 randomly chosen genes for these simulations.

One important metric to measure the performance of fine-mapping approaches is determining how well they identify implanted causal variants. For this measure, we define recall rate to be the proportion of genes in which all true causal variants are included in the causal set generated from the fine-mapping approaches. Since our method and the original CAVIAR method are designed to include all causal variants with 95% probability in their causal sets, their recall rate should be close to 95%. Simulation results show that our method has the correct recall rate for all numbers of causal variants and power levels (50% and 80%) for detecting a causal variant (Figure 2). The original CAVIAR method that uses only eQTL statistics achieves very similar recall rates as our method. These results demonstrate that our method detects all of true causal variants accurately even when there are several causal variants near a gene and when the power to detect a causal variant is not very high.

**Figure 2:**
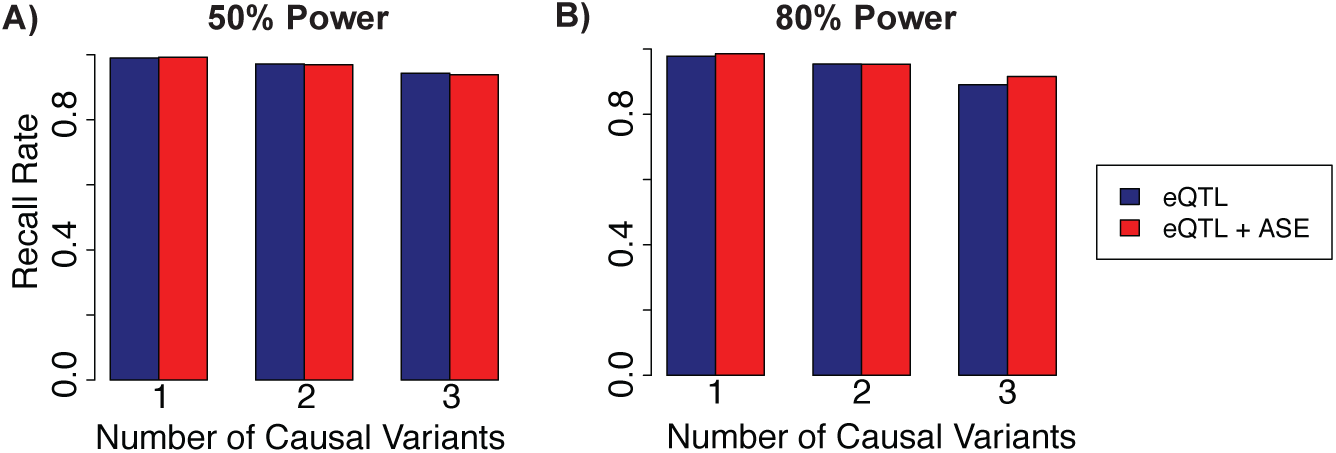
Recall rate comparison of our approach using meta-statistics (”eQTL+ASE”) and the previous approach using eQTL statistics (”eQTL”). Recall rate should be close to the designated 95% sensitivity level. Recall rate is measured for one, two, and three causal variants, and in two power levels; A) 50% power to detect causal variants and B) 80% power.

### 2.3 Meta-analysis reduces the size of causal set

A naive approach to improve recall rate would be to include as many variants as possible in the causal set. For instance, if we include all variants near a gene in the causal set, the recall rate would always be 1, as the causal set is guaranteed to contain all causal variants. This, however, would not be cost effective because downstream validation of these variants using functional assays is often costly. Therefore, the size of the causal set needs to be minimized while retaining high recall rate. While previous fine-mapping approaches such as CAVIAR attempt to minimize the causal set size, they may not accurately differentiate between causal and non-causal variants if they are in regions of high LD. Our method incorporates additional information about causal variants, ASE statistics, into our fine-mapping framework to improve our ability to determine whether variants are causal or not.

We compare the size of the causal set between our approach and the original CAVIAR approach in the following manner. For each gene, we calculate the *reduction rate* as 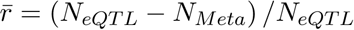 where *N*_*M eta*_ is the size of the causal set from our meta-analysis approach and *N*_*eQTL*_ is the size of the causal set from the original CAVIAR approach using only eQTL statistics. A positive reduction rate means our approach has the smaller causal set size than the original CAVIAR approach, while a negative reduction rate means it has the larger causal set. We compute the median reduction rate, which is the median of reduction rates among all genes.

Using the same simulation data as in the previous recall rate simulation, we show that the size of the causal set from our approach is noticeably smaller than that from the CAVIAR approach that uses only eQTL statistics (Figure 3). The median reduction rate is 24%, 31%, and 27% when the number of causal variants is 1, 2, and 3, respectively at 50% power to detect causal variants. At 80% power to detect causal variants, the median reduction rate is 27%, 31%, and 29% when the number of causal variants is 1, 2, and 3, respectively. This result also shows that we can achieve high reduction rates when there are multiple causal variants in a locus (Figure 3). This is advantageous because it is harder to predict causal variants when there are several causal variants that may have high LD with other non-causal variants. By incorporating information about causal variants for ASE, our framework is able to exclude non-causal variants from the causal set, reducing the causal set size.

**Figure 3:**
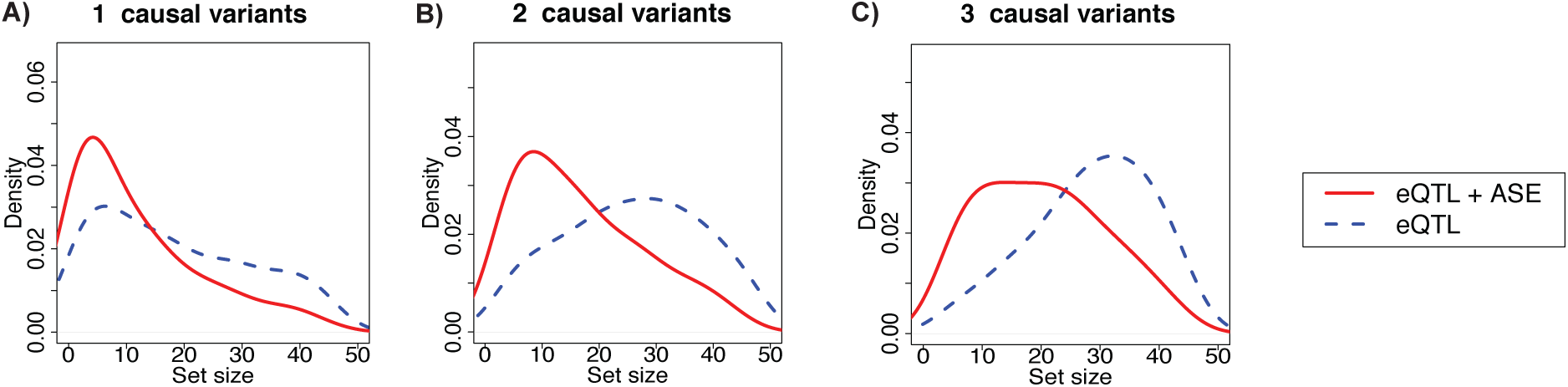
The distribution of causal set sizes in our simulation. We consider three scenarios where we have A) one causal variant, B) two causal variants, and C) three causal variants. The power to detect causal variants is 50% in all scenarios. “eQTL+ASE” shows the set size of our approach that uses meta-statistics while “eQTL” shows the set size of the previous approach using eQTL statistics.

### 2.4 Different effect sizes yield modest decrease in reduction rate

As our method utilizes fixed effect meta-analysis to combine eQTL and ASE statistics, it assumes that effect sizes in the eQTL studies and ASE studies are the same. To measure the performance of our approach when this assumption is violated, we consider scenarios where the true effect size in eQTL studies is not equal to that in ASE studies. Using the same 1000 randomly chosen genes from the previous simulation, we fix the effect size of the eQTL summary statistics to 5.2 (50% power) and vary the effect size of ASE summary statistics from 2.0 to 4.0. Then, using the same simulation framework, we generate ASE summary statistics and eQTL summary statistics. We observe that the median reduction rate decreases as the fixed effect assumption is further violated, which is expected (Figure 4A). The results also show that as long as ASE effect size is not prohibitively low compared to that of eQTL, our framework can yield a positive reduction rate; for all numbers of causal variants, the reduction rate is positive when the ASE effect size is greater than 2.0 (38% of eQTL effect size). Although we decrease the effect size of the ASE summary statistics in these simulations, the argument is symmetric, and decreasing the eQTL statistics should have similar results. In real data, although effect sizes of eQTL and ASE are unknown, it is very unlikely to observe such a large discrepancy between the two effect sizes. This also means that we expect slightly reduced reduction rate in real data compared to those observed in simulation (Figure 3) as there may be some genes in which effect sizes of eQTL and ASE are not exactly the same. However, in most cases, we expect to observe a positive reduction rate.

**Figure 4:**
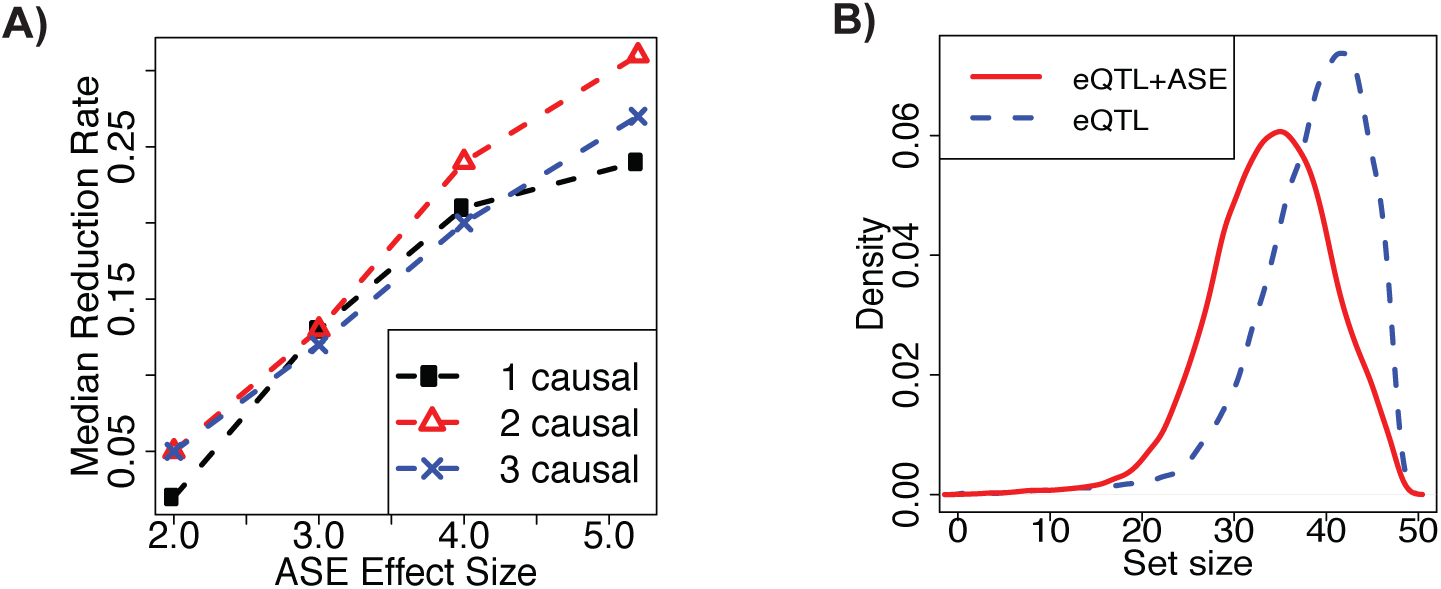
A) Median reduction rate using simulated data with different effect sizes between eQTL and ASE statistics. eQTL effect sizes are fixed at 5.2 while ASE effect sizes range from 2.0 to 4.0. B) The distribution of causal set sizes from the GTEx data for our approach using meta-statistics (”eQTL+ASE”) and the previous approach using only eQTL statistics (”eQTL”).

### 2.5 Meta-analysis improves fine-mapping in GTEx data

We then apply our approach to the RNA-seq data from GTEx [10]. We applied the framework to ten randomly selected tissues, ranging in sample sizes from 101 to 491. Unlike simulated data, real ASE data is not perfect, as it contains mapping errors and non-negligible noises that may reduce ASE signals significantly for certain genes. In particular, a small number of reads overlapping a certain gene may make it difficult to correctly identify individuals with ASE. This gene may contain very low ASE statistics for many genetic variants even though those genetic variants have high eQTL statistics. Our method may not behave correctly for this problematic gene, and hence we require individuals to have at least 20 reads mapped to each gene. While this requirement increases the quality of ASE calls for each individual, it may reduce the number of genes for which this method can be applied in tissues with small sample sizes (Table 1).

**Table 1:** Our method reduction rate for 10 GTEx tissues. Let (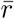) indicate the median reduction rate, *N* indicates the sample size, and *M* indicate number of genes.

| Tissue | $\bar{r}$ | N | M |
| --- | --- | --- | --- |
| Adipose Subcutaneous | 0.11 | 385 | 4023 |
| Artery Aorta | 0.11 | 267 | 2842 |
| Brain Caudate Basal Ganglia | 0.11 | 144 | 1272 |
| Brain Hippocampus | 0.10 | 111 | 680 |
| Heart Atrial Appendage | 0.11 | 264 | 1907 |
| Cells EBV-transformed Lymphocytes | 0.09 | 117 | 470 |
| Muscle Skeletal | 0.11 | 491 | 3414 |
| Thyroid | 0.10 | 399 | 1470 |
| Uterus | 0.09 | 101 | 600 |
| Whole Blood | 0.11 | 369 | 3452 |

We set the recall rate of our method and the original CAVIAR approach to 95%, meaning that both approaches generate causal sets that include all causal variants with 95% probability. Results show that our approach is effective at reducing the causal sets generated by the original CAVIAR approach. The median reduction rate across tissues is 11%, and the median reduction rate for each tissue ranges from 9% to 11% (Table 1). Our method achieves similar reduction rates for all tissues, even those with small sample sizes. Across tissues, the reduction rate is positive for 80% of genes, which indicates that our approach is able to reduce the causal sets for a majority of genes. The distribution of causal set size using the two sets of statistics is shown in Figure 4.

In this analysis, we identify ASE status of each individual by binarizing the ASE data with an empirically determined threshold (see Methods section). Instead of using the fixed threshold to determine the ASE status, we also binarized ASE using a binomial test in the thyroid tissue (see Methods section). When using the data from the binomial test in our fine-mapping framework, we observe a slight decrease in median reduction rate (from 10% to 9.4%). Furthermore, the number of genes for which the fine-mapping could be applied decreases from 1470 to 1344. This decrease is due to a decrease in the number of individuals with ASE for some genes. Therefore, we recommend using the empirically determined threshold to identify individuals with ASE.

## 3 Discussion

We developed a novel fine-mapping approach for eQTL studies that utilizes ASE information from RNA-seq data. Based on the insight that causal variants for total gene expression also influence gene expression in an allele-specific manner, we developed a new statistic that aggregates ASE and eQTL statistics through a fixed effect meta-analysis. We then incorporated the meta-analysis statistic into the fine-mapping algorithm for GWAS called CAVIAR, which takes into account LD structure among variants and multiple causal variants in a region. We used simulations to show that our approach achieves the correct recall rate and reduces the causal set size considerably compared to the fine-mapping approach that uses only eQTL statistics. We also show through simulations that our method is effective at reducing causal set size even when the fixed effect assumption is not met. When applying our method to the GTEx data set, we observed the median of 11% reduction rate in causal set size across the ten tissues studied. These results demonstrate that eQTL and ASE statistics may be integrated to improve fine-mapping and to effectively reduce the number of variants tested in downstream studies.

While this method is not the first to combine ASE and eQTL data to identify cis-regulatory variants, it is unique in that it uses ASE and eQTL data to perform fine-mapping while taking LD and multiple causal variants into account. The majority of methods that combine ASE and eQTL data focus on improving power to detect cis-regulatory variants rather than improving specificity of causal sets [33, 34]. Additionally, other methods compute a marginal likelihood for each variant instead of modeling the entire locus, making them unable to account for LD structure and multiple causal variants within each locus [33, 34, 35]. Therefore, these methods often rely on using a threshold to isolate a set of putative causal variants. They are suboptimal, as they can potentially identify many non-causal variants in LD with causal variants or fail to identify true causal variants [18, 36].

One key advantage of our approach is that we do not need to obtain additional data. Researchers can perform ASE calling on existing RNA-seq data to identify the ASE status of individuals for all genes and apply our approach using available total gene expression data and ASE calling data. Hence, our approach enables more accurate detection of causal variants regulating gene expression without the need for additional experiments.

One of the difficulties in identifying causal variants using ASE information is accurate identification of ASE calls from RNA-seq data. Previous studies have shown that noise caused by mapping errors, small numbers of reads, or inconsistencies between two variants used to call ASE in one locus can impact our ability to accurately detect ASE [37, 38, 32]. These problematic ASE calls may adversely affect our fine-mapping algorithm, and hence in addition to correcting for these when quantifying reads, we attempt to reduce them in our analysis by requiring individuals to have at least 20 reads mapped to a gene. We expect that ASE calling algorithms will improve in the near future, and better algorithms will enable more accurate identification of the ASE status of individuals and allow our method to perform fine-mapping in a greater number of genes.

## 4 Methods

### 4.1 Overview of CAVIAR generative model

Let *S* = [*s*_1_, *s*_1_, … *s*_*m*_] indicate the observed marginal statistics (e.g., z-scores) for a set of *m* variants, where *s*_*i*_ is the observed marginal statistic of *i*-th variant. We assume the computed marginal statistics follow an MVN distribution [18],

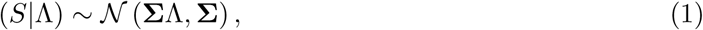

where **Σ** is the LD matrix between pairs of variants and Λ = [λ_1_, λ_2_, … λ_*m*_] is a vector of the true effect sizes [18]. Let *C* = [*c*_1_, *c*_1_, … *c*_*m*_] be a vector of zeros and ones that indicate the causal status of each variant. We define the prior probability on Λ for a given *C* as:

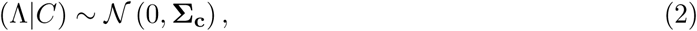

where **Σ**_c_ = *σ*^2^diag(C), and *σ* is a constant which indicates the variance of our prior over the true effect sizes. Similar to prior studies, we set *σ* to 5.2 [18, 22, 39].

The posterior predictive distribution (Equation 3) is also MVN, and we can use this distribution to compute the joint likelihood of the marginal statistics given a causal status.

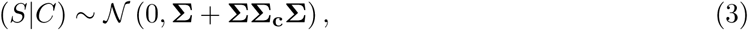

Given a set of possible causal statuses *ℂ*, the posterior probability of a causal status *C*^*^ ∈ *ℂ* can be expressed as 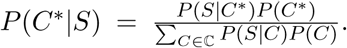. Using this fine-mapping framework, we can compute the posterior probability of causal status. Unfortunately, considering all possible causal sets is computationally intractable. To make this feasible, we assume that the maximum number of causal variants within a region is six [18, 22, 39]. We also use a greedy algorithm that eliminates the need to consider all possible subsets. To identify the minimal *ρ* causal set, at each iteration of the greedy algorithm, we select the variant that increases the total posterior probability the most. Variants are added until the posterior probability of the causal set is at least *ρ* fraction of the total posterior probability of the data.

### 4.2 Computing eQTL association statistics

Let *Y* be the normalized total expression values for *n* individuals in a single gene, and let *X*_*i*_ be the normalized genotypes of variant *i* for all individuals. Suppose *Y* = *μ* + *β*_*i*_*X*_*i*_ + ∈, where *μ* is the phenotypic mean in the population, *β*_*i*_ is the effect size of variant *i*, and ∈ models environmental and measurement noise. We use the maximum likelihood estimates 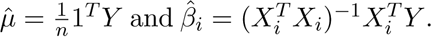 The error can be calculated as 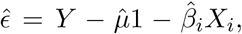 and the standard deviation is calculated as 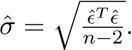 Using these, the association statistic for variant *i* is calculated as 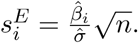

Assuming that we have enough individuals, the marginal statistics have the same posterior predictive distribution used by CAVIAR (Equation 3). The distribution of the eQTL summary statistics given a causal status is as follows:

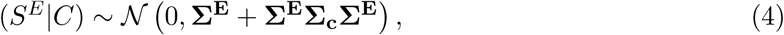

where *S*^*E*^ is a vector of the observed marginal statistics for a gene and **Σ**^**E**^ is the LD matrix between pairs of variants in the eQTL study.

### 4.3 Computing ASE association statistics

Using phased genotype data, ASE status for a gene in an individual can be directly computed from RNA-seq data by using heterozygous coding SNPs to map reads to one of the haplotypes. The proportion of reads mapping to each haplotype can be used as a proxy for relative contribution from each haplotype to total expression. The read mapping was performed by the GTEx consortium and accounts for genotyping error, reference bias, and other sources of technical variation [32]. Let *c*_1_ and *c*_2_ indicate the number of reads supporting the first and second haplotype, respectively. We calculate the allelic ratio (AR) for an individual as AR = 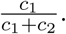 We consider an individual to have ASE for a gene when the allelic ratio is less than .35 or greater than .65; otherwise, we label the gene in the individual as having balanced expression. This threshold was obtained empirically in [31]. We compare this binarization framework to one using a binomial test for each individual. If the allelic ratio for an individual is significantly different from 0.5, the individual is labeled as having ASE. Otherwise, the individual is labeled as having balanced expression. The p-values for each individual are adjusted for multiple hypothesis testing using the Benjamini-Hochberg procedure.

If an individual is heterozygous for a causal variant, we expect the expression from each allele to be different. On the other hand, if an individual is homozygous for a causal variant, we expect the expression for each allele to be comparable. ASE association statistics measure the correlation between ASE status (e.g., 0 for balance expression and 1 for allelic imbalance) and heterozygous status of a variant (e.g. 0 for homozygous genotype and 1 for heterozygous genotype).

The association statistics are calculated in a way similar to case and control GWAS association statistics. Let 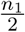 be the number of individuals with ASE in the study, and let 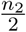 be the number of individuals with balanced expression in the study. Let *p*_1_ be the proportion of individuals with ASE who are heterozygous for the SNP. Let *p*_2_ be the proportion of individuals with balanced expression who are heterozygous for the SNP. The difference between these two proportions is also normally distributed. Under the null hypothesis (*p*_1_ = *p*_2_), we have:

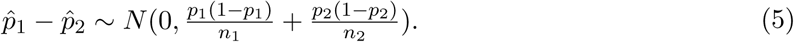

Let *p* be the frequency of heterozygous individuals in the population. With the simplifying assumption used in standard GWAS with unequal case and control size that 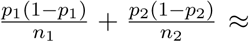 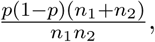 the null hypothesis becomes 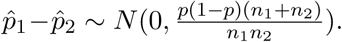 We set the ASE association statistic to be:

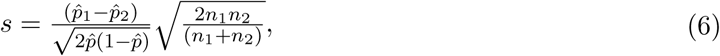

which follows the standard normal distribution.

We assume that the ASE summary statistics follow MVN, and the marginal statistics have the same posterior predictive distribution used by CAVIAR. We compute the heterozygosity matrix from genotype data for all variants in that locus. Thus, similar to the posterior predictive of CAVIAR (Equation 3) we have:

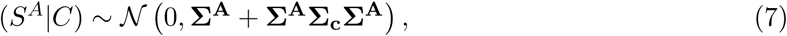

where *S*^*A*^ is the observed marginal statistics for the gene we are interested and **Σ**^**A**^ is the heterozygosity matrix.

### 4.4 Utilizing fixed-effect meta-analysis

Since we assume that ASE and eQTL studies have the same causal variants, we perform a fixed effect meta-analysis. The computed combined marginal statistics *S*^*M*^ can be calculated as 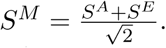 This combined statistic is the sum of two Gaussian random variables. Therefore, the posterior predictive can be expressed as follows:

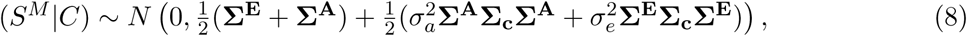

Since it is possible for ASE and eQTL summary statistics to be in different directions in the real data, we calculate two meta statistics, 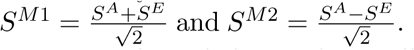 We perform fine-mapping using each set of meta-analysis statistics separately and choose the smallest causal set.

### 4.5 Calculating pairwise correlation matrices

To calculate pairwise correlation matrices, we use genotype data from the whole blood tissue in GTEx (Release v6, dbGaP Accession phs000424.v6.p1 available at: www.gtexportal.org) that contains 325 samples. For each gene, we compute the eQTL statistic for every *cis* variant within 1MB from the transcription start site and identify the top 50 genetic variants with the highest eQTL statistics. We use these top 50 genetic variants to compute the pairwise correlation matrices. For eQTL statistics, we calculated **Σ**^**E**^ as the pairwise correlation matrices between genotypes of individuals. For ASE statistics, we calculated **Σ**^**A**^ as the pairwise correlation between the heterozygosity calls of individuals. Since the statistics are normally distributed, we calculated the meta-analysis correlation matrix as 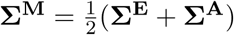

### 4.6 Generating simulated datasets

Given a genotype LD matrix (**Σ**^**E**^) and a heterozygosity LD matrix (**Σ**^**A**^) for a gene and a vector C that indicates the causal status of each SNP (e.g., 0 when the variant is not causal and 1 when the variant is causal), we can simulate the association statistics by sampling 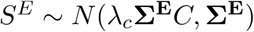) and 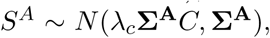, where *λ*_*c*_ is the non-centrality parameter (NCP). We set *λ*_c_ such that we have the desired power. We first set the power level to 50% and 80% for both ASE and eQTL statistics. Simulations at these power levels indicate that our fine-mapping method is effective for the strongest eQTL loci. We then explore a larger range of power in Section 2.4 and show that our method can also reduce causal set size in loci with lower power. We calculate the simulated statistics for the meta-analysis in two ways: 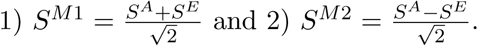. We perform one simulation for each gene in the GTEx data to observe a natural range of correlation between variants. Simulations to generate *S*^*E*^, *S*^*A*^, *S*^*M1*^, and *S*^*M2*^ were performed for up to three causal variants.

### 4.7 Application to GTEx data

To measure the total expression level of each gene, RNA-seq data was aligned to hg19 using Tophat v1.4.1 [40] and gene-level expression quantification was performed using RNA-SeQC [41] according to the GTEx protocol [8]. Reads were assigned to each allele of an individual using personalized genomes and the official GTEx protocol described in [32]. To reduce the affect of noise in ASE calls, we require individuals to have at least 20 reads mapped to each gene. For genotype data, we used the Illumina OMNI 5M SNP Array. To increase eQTL discovery power, we used genotypes that were imputed from the 1000 Genomes Project Phase I version 3 reference panel using the IMPUTE2 software [42].

We apply this framework to genes with at least one significant eQTL identified in the GTEx data set and at least one heterozygous variant in the coding region. The significant eQTL requirement restricts the set of genes tested to those likely to be regulated by cis-regulatory variants. The heterozygous variant in the coding region is used to call ASE. For each gene, we selected the top 50 variants that have the highest eQTL statistics. We calculate the meta statistics and heterozygosity matrix for these variants, which we use as input to our fine-mapping framework. In some cases, the eQTL statistic and the ASE statistic have different signs. In these cases, the statistics cancel each other out in *S*^*M1*^. Therefore, we calculate *S*^*M2*^, and run CAVIAR on both sets of summary statistics. We take the minimum of these two data sets.

## Acknowledgment

J.Z., F.H., and E.E. are supported by National Science Foundation grants 0513612, 0731455, 0729049, 0916676, 1065276, 1302448, 1320589 and 1331176, and National Institutes of Health grants K25-HL080079, U01-DA024417, P01-HL30568, P01-HL28481, R01-GM083198, R01-ES021801, R01-MH101782 and R01-ES022282. E. E. is supported in part by the NIH BD2K award, U54EB020403. J.Z and J.E. are supported by National Institute of Health grants R01ES024995, U01HG007912, DP1DA044371, NSF CAREER Award #1254200, and an Alfred P. Sloan Fellowship (J.E.). J.Z. is supported by National Institutes of Health Award Number T32MH073526. We acknowledge the support of the NINDS Informatics Center for Neurogenetics and Neurogenomics (P30 NS062691). The authors declare that they have no competing interests.

## References

[1] Valur Emilsson, et al. Genetics of gene expression and its effect on disease. Nature, 452(March), 2008.

[2] Alexandra C Nica, et al. Candidate causal regulatory effects by integration of expression qtls with complex trait genetic associations. PLoS Genetics, 6(4):e1000895, 2010.

[3] Dan L Nicolae, et al. Trait-associated snps are more likely to be eqtls: annotation to enhance discovery from gwas. PLoS Genetics, 6(4):e1000888, 2010.

[4] Lea K. Davis, et al. Partitioning the heritability of tourette syndrome and obsessive compulsive disorder reveals differences in genetic architecture. PLoS Genetics, 9(10):e1003864, oct 2013.

[5] Jason M. Torres, et al. Cross-tissue and tissue-specific eQTLs: Partitioning the heritability of a complex trait. The American Journal of Human Genetics, 95(5):521–534, nov 2014.

[6] Rachel B Brem and Rebecca Clinton. Genetic Dissection of Transcriptional Regulation in Budding Yeast. Science, 296(April), 2002.

[7] John Lonsdale, et al. The Genotype-Tissue Expression (GTEx) project. Nat Genet, 45(6):580– 585, jun 2013.

[8] Kristin G Ardlie, et al. The genotype-tissue expression (gtex) pilot analysis: Multitissue gene regulation in humans. Science, 348(6235):648–660, 2015.

[9] Daria V Zhernakova, et al. Identification of context-dependent expression quantitative trait loci in whole blood. Nature Genetics, 49(1):139–145, dec 2016.

[10] GTEx Consortium, et al. Genetic effects on gene expression across human tissues. Nature (accepted), 2017.

[11] The Wellcome, et al. Articles Bayesian refinement of association signals for 14 loci in 3 common diseases. Nature Genetics, 44(12), 2012.

[12] Nathalie Malo, et al. Accommodating Linkage Disequilibrium in Genetic-Association Analyses via Ridge Regression. Am. J. Hum. Genet., 82(February):375–385, 2008.

[13] Jian Yang, et al. Conditional and joint multiple-SNP analysis of GWAS summary statistics identifies additional variants influencing complex traits. Nature Publishing Group, 44(4):369– 375, 2012.

[14] Rick Jansen, et al. Conditional eQTL analysis reveals allelic heterogeneity of gene expression. Human Molecular Genetics, feb 2017.

[15] Farhad Hormozdiari, et al. Widespread allelic heterogeneity in complex traits. The American Journal of Human Genetics, 2017 (In press).

[16] A. Andrew Brown, et al. Predicting causal variants affecting expression by using whole-genome sequencing and RNA-seq from multiple human tissues. Nature Genetics, Oct 2017.

[17] Bertrand Servin and Matthew Stephens. Imputation-based analysis of association studies: Candidate regions and quantitative traits. PLoS Genetics, 3(7):1296–1308, 2007.

[18] Farhad Hormozdiari, et al. Identifying causal variants at loci with multiple signals of association. Genetics, 198(2):497–508, 2014.

[19] Kyle Kai-How Farh, et al. Genetic and epigenetic fine mapping of causal autoimmune disease variants. Nature, 518(7539):337–343, oct 2014.

[20] Wenan Chen, et al. Fine mapping causal variants with an approximate bayesian method using marginal test statistics. Genetics, 200(3):719–736, 2015.

[21] Christian Benner, et al. FINEMAP: efficient variable selection using summary data from genome-wide association studies. Bioinformatics, 32(10):1493–1501, 2016.

[22] Farhad Hormozdiari, et al. Identification of causal genes for complex traits. Bioinformatics, 31(12):i206–i213, 2015.

[23] Tomi Pastinen and Thomas J Hudson. Cis-acting regulatory variation in the human genome. Science (New York, N.Y.), 306(5696):647–650, 2004.

[24] Yehudit Hasin-Brumshtein, et al. Allele-specific expression and eqtl analysis in mouse adipose tissue. BMC Genomics, 15(1):471, Jun 2014.

[25] Y. Baran, et al. The landscape of genomic imprinting across diverse adult human tissues. Genome Research, 25(7):927–936, Jul 2015.

[26] Pejman Mohammadi, et al. Quantifying the regulatory effect size of cis-acting genetic variation using allelic fold change. Genome Research, pages 1–13, 2017.

[27] H. Yan. Allelic Variation in Human Gene Expression. Science, 297(5584):1143–1143, aug 2002.

[28] Dominique J. Verlaan, et al. Targeted screening of cis-regulatory variation in human haplotypes. Genome Research, pages 118–127, 2009.

[29] Tomi Pastinen. Genome-wide allele-specific analysis: insights into regulatory variation. Nature Reviews Genetics, 11(8):533–538, aug 2010.

[30] Kun Zhang, et al. Digital RNA allelotyping reveals tissue-specific and allele-specific gene expression in human. Nature methods, 6(8):613–618, 2009.

[31] Eun Yong Kang, et al. Discoving SNPs Regulating Human Gene Expression Using Allele Specific Expression from RNA-Seq data. Genetics, 204:1–12, 2016.

[32] Matti Pirinen, et al. Assessing allele-specific expression across multiple tissues from RNA-seq read data. Bioinformatics, 31(15):2497–2504, 2015.

[33] Bryce van de Geijn, et al. WASP: allele-specific software for robust molecular quantitative trait locus discovery. Nature Methods, 12(11):1061–3, 2015.

[34] Y Hu, et al. Proper Use of Allele-Specific Expression Improves Statistical Power for cis-eQTL Mapping with RNA-Seq Data. J Am Stat Assoc, 110(511):962–974, 2015.

[35] Natsuhiko Kumasaka, et al. technical reports Fine-mapping cellular QTLs with RASQUAL and ATAC-seq. Nature Genetics, 48(2), 2016.

[36] Miriam S Udler, et al. FGFR2 variants and breast cancer risk: fine-scale mapping using African American studies and analysis of chromatin conformation. Human Molecular Genetics, 18(9):1692–1703, 2009.

[37] Chris T Harvey, et al. Genetics and population analysis QuASAR: quantitative allele-specific analysis of reads. Bioinformatics, 31(December 2014):1235–1242, 2015.

[38] Jacob F. Degner, et al. Effect of read-mapping biases on detecting allele-specific expression from RNA-sequencing data. Bioinformatics, 25(24):3207–3212, dec 2009.

[39] Farhad Hormozdiari, et al. Colocalization of GWAS and eQTL signals detects target genes. The American Journal of Human Genetics, 99(6):1245–1260, dec 2016.

[40] Cole Trapnell, et al. TopHat: discovering splice junctions with RNA-Seq. Bioinformatics, 25(9):1105–1111, 2009.

[41] DS DeLuca, et al. Rna-seqc: Rna-seq metrics for quality control and process optimization. Bioinformatics, 11(28):1530–2, 2012.

[42] Bryan N Howie, et al. A Flexible and Accurate Genotype Imputation Method for the Next Generation of Genome-Wide Association Studies. PLoS Genetics, 5(6), 2009.

